# Progressive chromatin silencing of ABA biosynthesis gene permits seed germination in *Arabidopsis*

**DOI:** 10.1101/2021.10.08.463712

**Authors:** Deyue Yang, Fengli Zhao, Danling Zhu, Xi Chen, Xiangxiong Kong, Yufeng Wu, Min Chen, Jiamu Du, Li-jia Qu, Zhe Wu

## Abstract

Seed germination represents a major developmental switch in plants that is vital to agricultural, but how this process is controlled at the chromatin level remains obscure. Here we demonstrate that successful germination in *Arabidopsis* requires a chromatin mechanism that progressively silences *NCED6*, which encodes a rate-limiting enzyme for abscisic acid (ABA) biosynthesis, through the cooperative action of the RNA-binding protein RZ-1 and the polycomb repressive complex 2 (PRC2). Simultaneous inactivation of RZ-1 and PRC2 blocks germination and synergistically derepresses *NCEDs* and hundreds of genes. At *NCED6*, by promoting H3 deacetylation and suppressing H3K4me3, RZ-1 facilitates transcriptional silencing and also a H3K27me3 accumulation process that occurs during seed germination and early seedling growth. Genome-wide analysis reveals RZ-1 is preferentially required for transcriptional silencing of many PRC2 targets early during seed germination when H3K27me3 is not yet established. We propose RZ-1 confers a novel silencing mechanism to compensate and coordinate with PRC2. Our work highlights the progressive chromatin silencing of ABA biosynthesis genes via synergized action of the RNA-binding protein RZ1 and PRC2, which is vital for seed germination.

## Introduction

Plant seed is a unique form of life reservoir on earth (Linkies et al., 2010). In the dry seeds of many plants like *Arabidopsis thaliana*, cellular activities such as transcription cease, allowing the seed to resist harsh challenges from the environment. Strikingly, under favourable conditions and upon water uptake (imbibition), the seed quickly resumes its cellular activities, germinates and develops into a vulnerable seedling within only a few days, representing one of the most critical developmental switches in flowering plants (Finch-Savage and Leubner-Metzger, 2006).

Any successful developmental switch relies on the reprograming of gene expression. Whereas some genes are activated by pioneer transcription factors (Iwafuchi-Doi and Zaret, 2014; Zaret and Mango, 2016), many others are progressively switched off and maintained in a silenced state epigenetically, forming facultative heterochromatin (Berry and Dean, 2015; Brockdorff et al., 2020). Although the roles of transcription factors in seed germination have been studied in-depth, little is known about how seed germination operates at the chromatin level. The inactivation of polycomb repressive complex 2 (PRC2) or PRC1 leads to delayed seed germination and the ectopic expression of seed-specific genes (e.g., *ABI3*, *DOG1*) in seedlings, suggesting that these complexes function in the chromatin silencing of specific genes at the onset of seed-to-seedling transition (Bouyer et al., 2011; Yang et al., 2013; Molitor et al., 2014). However, the importance of such epigenetic repression at individual genes for seed germination *per se* is unknown. Interestingly, although plant PRC2 is essential for proper development post germination, it is not indispensable for embryo development and seed germination(Bouyer et al., 2011). Indeed, only a subset of H3K27me3 marked genes are strongly derepressed in *swn clf* mutant (Bouyer et al., 2011), suggesting the presence of unknown silencing mechanism that compensate for PRC2 at its targets (Bouyer et al., 2011). To date, the nature of such silencing mechanism is unknown.

The phytohormone abscisic acid (ABA) plays a fundamental role in seed germination. Artificially inducing expression of the rate-limiting ABA biosynthesis enzyme nine-*cis*-epoxycarotenoid dioxygenase 6 (NCED6) or NCED9 strongly delays germination (Martinez-Andujar et al., 2011), suggesting that restricting ABA biosynthesis is necessary for germination. In newly harvested seeds with primary dormancy, both *NCED6* and *NCED9* can be activated by transcription factor MYB96 (Lee et al., 2015) and the basic helix-loop-helix transcription factor bHLH57 (Liu et al., 2020). During seed germination, *NCED9* can be activated by overexpressing the transcription factor DEHYDRATION-RESPONSIVE ELEMENT BINDINGFACTOR2C (DREB2C) (Je et al., 2014), while *NCED6* can be activated by the loss of function of ABSCISIC ACID INSENSITIVE4 (ABI4) (Barros-Galvao et al., 2020). To date, for a non-dormant seed, although the ABA biosynthesis is known to be suppressed during its germination, the mechanism through which the suppression is achieved is largely unknown.

Previously, we characterized two functionally redundant RNA-binding proteins, RZ-1B and RZ-1C (RZ-1) in *Arabidopsis* (Wu et al., 2016b). Loss of RZ-1 leads to pleiotropic phenotypes including delayed seed germination, accompanied with disrupted expression of hundreds of genes (Wu et al., 2016b). RZ-1 associates with chromatin (Wu et al., 2016b) and is required for efficient co-transcriptional splicing (Zhu et al., 2020), an activity that associates with its direct binding to nascent RNAs (Zhu et al., 2020).

Here, we report an RZ-1 mediated chromatin silencing mechanism that is crucial for seed germination. We show that RZ-1 promotes seed germination by mediating a transcriptional silencing mechanism that tightly cooperates with PRC2 mediated silencing, including at *NCED6*, a gene responsible for the delayed germination of *rz-1.* RZ-1 is required for active histone deacetylation and suppression of H3K4me3 by interacting with histone deacetylase and H3K4me3 reader proteins. This process leads to transcriptional silencing and also enhances the timely establishment of H3K27me3 marks mediated by PRC2. Intriguingly, RZ-1 is mainly required for transcriptional silencing of PRC2 targets when H3K27me3 has not been established yet during seed germination. Our work thus reveals a silencing mechanism mediated by RNA-binding protein RZ-1 which compensates for and coordinates with PRC2 mediated silencing, which is vital for seed germination.

## Results

### The absence of both RZ-1B and RZ-1C delays seed germination and alters sensitivity to an exogenous ABA biosynthesis inhibitor in *Arabidopsis*

To dissect the mechanism underlying delayed seed germination in *rz-1b/1c*, we investigated whether this phenotype is related to the ABA or gibberellic acid (GA) pathway by examining the germination rates of after-ripened seeds of Col-0, *rz-1b rz-1c* and *GFP-RZ-1C* (*rz-1b rz-1c* mutant complemented with a GFP-RZ-1C fusion driven by its own promoter and genomic fragment (Wu et al., 2016b)) in the presence of these hormones or their biosynthesis inhibitors. In the absence of hormones (Figure 1a, b), *rz-1b rz-1c* showed delayed germination (∼36 h delay in reaching a 50% germination rate) compared to Col-0 or the complemented line. GA treatment had a subtle effect on the germination of *rz-1b rz-1c* (Supplementary Figure 1), indicating that the delayed germination of *rz-1b rz-1c* is not due to a shortage of GA. By contrast, ABA treatment exaggerated the delayed germination of *rz-1b rz-1c*, whereas treatment with norflurazon (an ABA biosynthesis inhibitor) markedly accelerated seed germination in *rz-1b rz-1c* but not in Col-0 or *GFP-RZ-1C* (Figure 1c). Thus, altered ABA biosynthesis, decay or signalling likely contributes to the delayed germination in *rz-1b rz-1c*.

**Figure 1.**
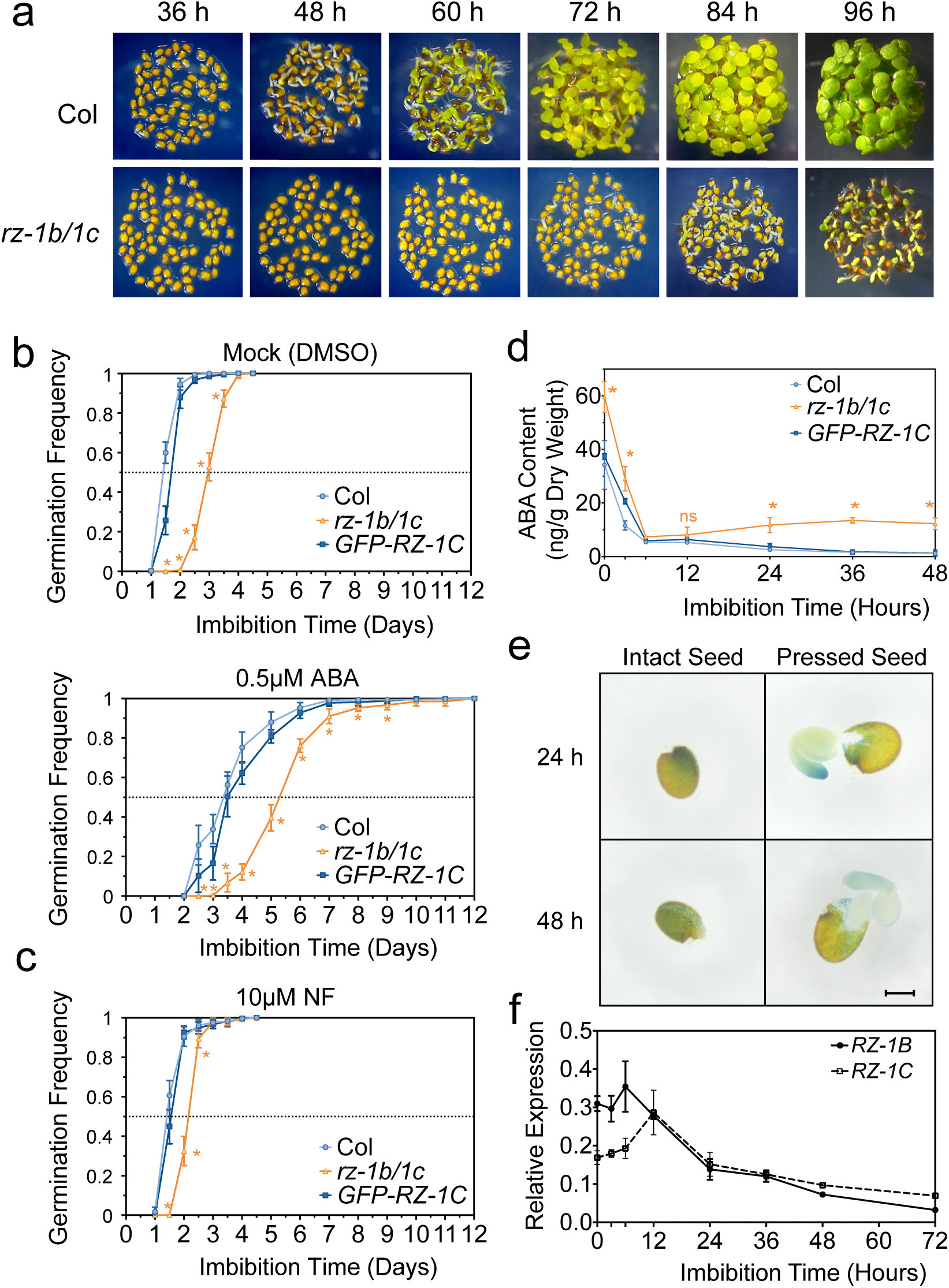
The absence of RZ-1 leads to delayed germination due to elevated ABA content. **a)** Representative photographs of Col-0 and *rz-1b/1c* during seed imbibition and early seedling development on ½ MS medium. **b and c)** Statistics of germination frequency over a time course of imbibition. Seeds were sown directly on ½ MS medium supplemented with solvent (DMSO) as a mock control (b), 0.5 μM abscisic acid (ABA) (b) or 10 μM of the ABA biosynthesis inhibitor norflurazon (NF) (c). Data are presented as mean ± SD from six biological replicates, with 50 seeds per replicate. The horizontal dotted line indicates 50% germination frequency. **d)** ABA contents in dry and imbibed seeds of Col-0, *rz-1b/1c* and GFP-RZ-1C *rz-1b/1c* over a time course. Data are presented as mean ± SD from three biological replicates. For b to d, * indicates significant difference between Col-0 and *rz-1b/1c* based on two-tailed Student’s *t*-test at *p* < 0.05. **e)** Representative photographs of GUS staining of seeds of the *rz-1b/1c* double mutant complemented with the *RZ-1C: GUS-GFP-RZ-1C* reporter construct at 24 h or 48 h post-imbibition. ‘Pressed’ seeds were obtained by poking ‘intact’ seeds gently to separate the embryo from the seed coat and endosperm. Scale bar indicates 0.2mm. **f)** Time course of mRNA expression of *RZ-1B* and *RZ-1C* during seed imbibition. Values were normalized to *UBC9* as a reference, and data are presented as mean ± SD from three biological replicates.

### *rz-1b rz-1c* shows increased ABA contents during seed imbibition

To determine whether ABA level is increased in *rz-1b rz-1c*, we measured ABA contents in dry and imbibed seeds of different genotypes over a time course (Figure 1d). In dry seeds, the ABA level was substantially higher in *rz-1b rz-1c* compared to Col-0 and *GFP-RZ-1C*. Upon imbibition, however, all genotypes showed rapid declines in ABA content and reached similar levels by 6 h, suggesting that ABA catabolism underlying this process is intact in *rz-1b rz-1c.* Importantly, from 12 h to 48 h of imbibition, an increase in ABA content was clearly observed in *rz-1b rz-1c*. By contrast, in Col-0 and *GFP-RZ-1C,* the ABA contents continued to decline throughout the entire imbibition period. Consistently, RZ-1C protein is expressed in tissues where ABA is synthesized (Lefebvre et al., 2006), including the embryo and endosperm, during seed germination (Figure 1e), as revealed by histochemical staining using a transgenic line expressing *GUS-GFP-RZ-1C* driven by the native *RZ-1C* promoter; this construct was also able to complement the *rz-1b rz-1c* phenotype. In addition, qPCR analysis showed that *RZ-1B* and *1C* transcript levels peaked within the first 12 h of imbibition and declined gradually thereafter (Figure 1f). These data suggest that RZ-1B and 1C function redundantly and actively during seed germination to limit ABA levels in seeds.

### The loss function of RZ-1 disrupts the progressive silencing of ABA biosynthesis genes during germination

To investigate the cause for increased ABA accumulation in *rz-1b/1c*, we performed mRNA-seq using 24 h imbibed seeds. As expected, many ABA pathway genes were expressed at higher levels in *rz-1b/1c* than in the wild type (Figure 2a). Notably, *NCED2/4/5/6/9*, which encode rate-limiting enzymes of ABA biosynthesis, were upregulated in *rz-1b/1c* (Figure 2a). Next, we measured the expression of these genes over a 72-hour time course upon imbibition, and at 6 days, 10 days and 15 days post imbibition (Figure 2b). *NCED2/5/6/9* became progressively silenced during germination in an RZ-1-dependent manner. Specifically, shortly after imbibition (∼6 h), the expression levels of *NCEDs* in Col-0 and *GFP-RZ-1C* declined rapidly and remained at a repressed state thereafter. By contrast, in *rz-1b/1c*, expression of these genes was upregulated to various extents after 12 or 24 h of imbibition and remained at relatively high levels for the remainder of imbibition (Figure 2b). Notably, *NCED6* was expressed at several hundred-fold higher levels in *rz-1b/1c* than that in Col-0 from 36 h to 72 h. In addition, ABA signalling genes *ABI3* and *ABI5* were also repressed by RZ-1 (Figure 2c). By contrast, ABA catabolism genes were much less affected by RZ-1 (Figure 2d). In seedlings (6 days, 10 days and 15 days), *NCEDs* were generally repressed even in *rz-1b/1c*, although they were still at higher levels than that in Col-0 (e.g., *NCED6*) (Figure 2b). Therefore, additional factors are expected to contribute to the silencing program of *NCEDs* during the seed-to-seedling switch. In conclusion, RZ-1 is required for a mechanism that operates during germination to progressively silence *NCEDs*.

**Figure 2.**
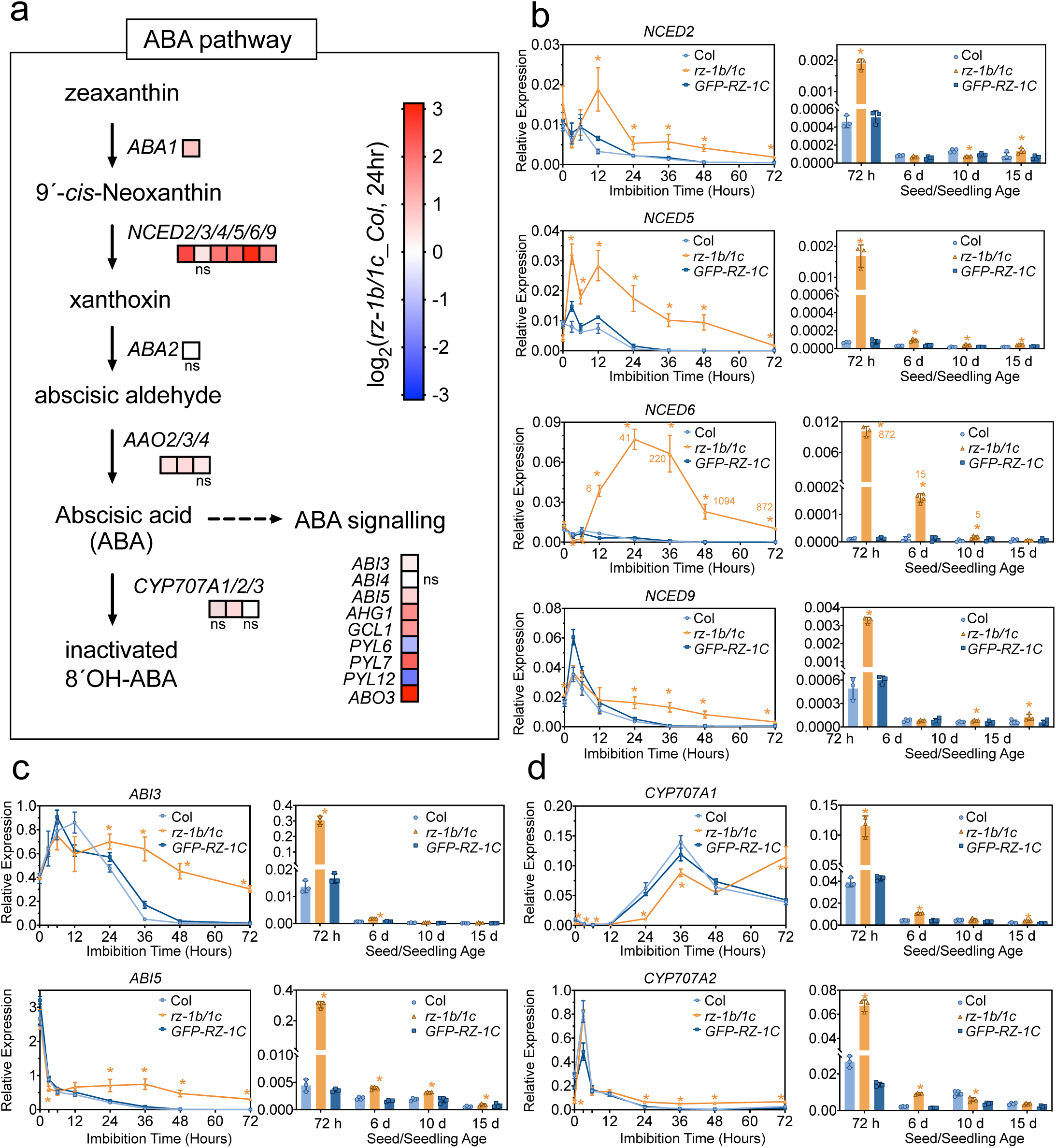
*rz-1b/1c* disrupts the progressive silencing of ABA biosynthesis genes during germination. **a)** Illustration of the ABA metabolic pathway with altered gene expression in *rz-1b/1c* compared to Col-0 after 24-h imbibition based on mRNA-seq data. **b–d)** qPCR analysis of the expression of ABA pathway genes in Col-0, *rz-1b/1c* and *GFP-RZ-1C rz-1b/1c* over an imbibition time course; b) ABA biosynthesis genes *NCED2/5/6/9*; c) ABA signaling genes *ABI3* and *ABI5*; d) ABA catabolism genes *CYP707A1* and *CYP707A2*. Values were normalized to *UBC9* and are presented as mean ± SD from at least three biological replicates. Individual data points are also indicated in bar charts and 72 h data are shown in both diagrams to aid visualization. * indicates significant difference between Col-0 and *rz-1b/1c* based on two-tailed Student’s *t*-test at *p* < 0.05. For *NCED6*, relative expression (fold change) in *rz-1b/1c* compared to Col-0 is indicated by a number adjacent to the mean at each time point.

### Upregulation of ABA biosynthesis through *NCED6* is mainly responsible for delayed germination in *rz-1b rz-1c*

To test whether the disrupted silencing of *NCEDs* is responsible for the delayed germination in *rz-1b/1c*, we crossed *rz-1b/1c* with ABA-related mutants to generate respective triple mutants. Germination analysis revealed almost full recovery in *aba1 rz-1b/1c* (Figure 3a, b), indicating RZ-1 controls germination through ABA pathway, given that ABA1 functions as the only enzyme in the first step of ABA metabolism. A similar recovery in germination was also observed in the *nced6 rz-1b/1c* triple mutant (Figure 3a, b), whereas crossing with other *nced* mutants barely recovered germination in *rz-1b/1c* (Figure 3b), indicating that the upregulation of *NCED6,* but not other *NCEDs,* is mainly responsible for the delayed germination in *rz-1b/1c*. Additionally, the delayed germination was also partially restored in *abi3 rz-1b/1c* triple mutant (Figure 3a, b), which is consistent with ABI3’s role as a key ABA signalling factor in seeds. Notably, although there is very low level of *ABI3* transcribed from the *abi3* locus, it is unlikely responsible for the incomplete restoration of seed germination in *abi3 rz-1b/1c* (Supplementary Figure 2). Therefore, the upregulated ABA biosynthesis due to derepression of *NCED6* is mainly responsible for the delayed germination in *rz-1b/1c*.

**Figure 3.**
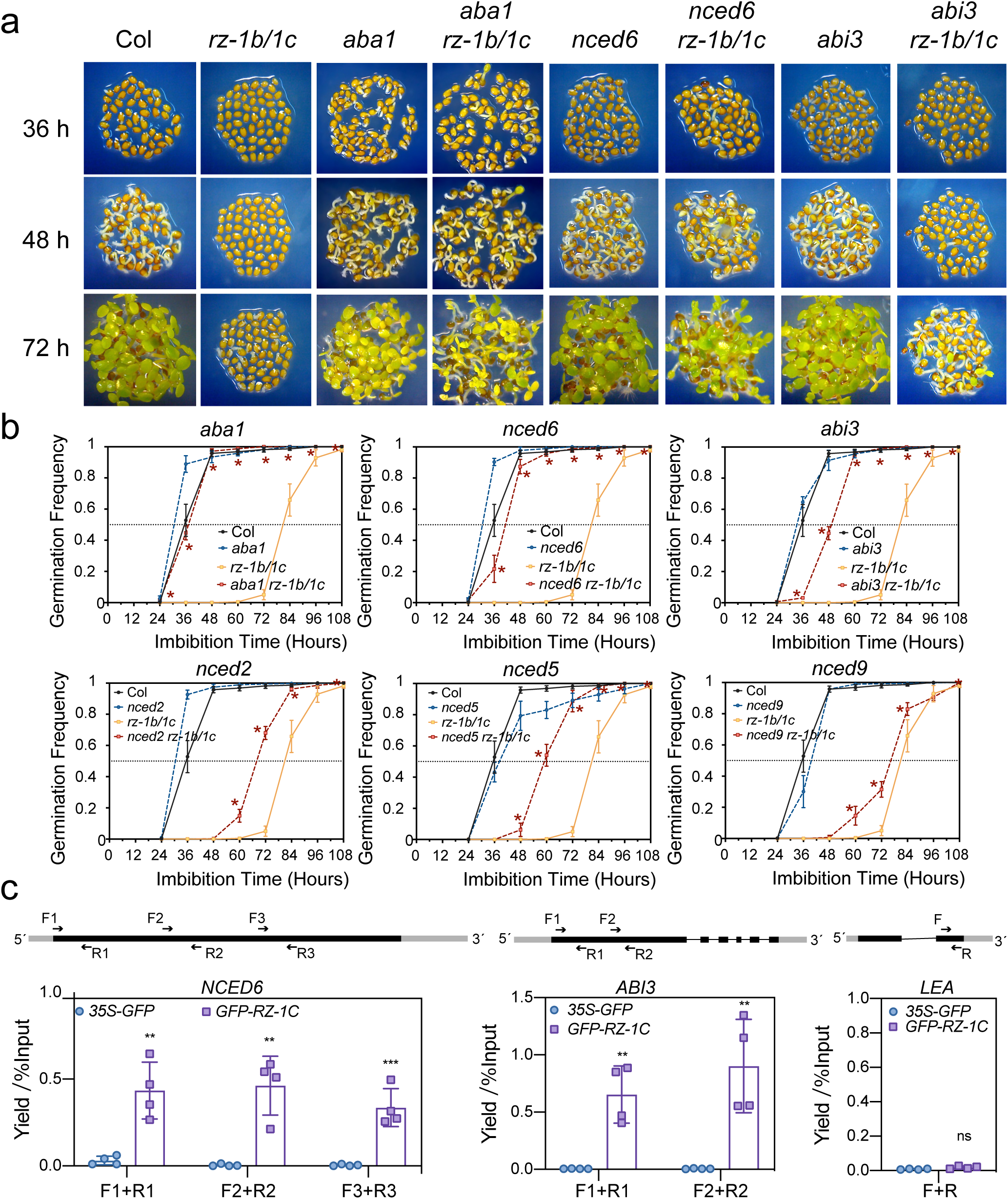
Upregulation of ABA biosynthesis through *NCED6* is mainly responsible for delayed germination in *rz-1b rz-1c*. **a)** Representative photographs of Col-0, *rz-1b/1c*, ABA-deficient mutants and triple mutants generated by genetic crossing during seed germination. **b)** Germination frequency analyses of various ABA mutants and each respective crossing with *rz-1b/1c*. Data are presented as mean ± SD from five biological replicates, with 50 seeds per replicate. * indicates significant difference between *rz-1b/1c* and the corresponding triple mutant based on two-tailed Student’s *t*-test at *p* < 0.05. **c)** RIP-qPCR showing the specific binding of RZ-1C to nascent RNAs of *NCED6* and *ABI3*; *LEA* was used as a negative control without RZ-1C binding. Gene structures are illustrated using black boxes as exon, single lines as introns and grey boxes as UTRs. Locations of primers are indicated using lines with arrows. Data were normalized to 1% of input and are presented as mean ± SD from four biological replicates. ** and *** indicate statistical significance based on Student’s *t*-test at *p*-values < 0.01 and 0.001, respectively. ns = not significant.

### *NCED6* and *ABI3* are binding targets of RZ-1C at chromatin level

We previously demonstrated that RZ-1C is a chromatin associated protein that binds RNA in vitro and in vivo (Wu et al., 2016b). We therefore tested whether *NCED6* is a target of RZ-1 at the chromatin associated RNA level. Unfortunately, our attempts at unbiased mapping of the RNA-binding targets of RZ-1C in seeds by eCLIP-seq were unsuccessful, likely due to the blocking of UV light penetrance by the rigid layer of the seed coat. We therefore performed a formaldehyde crosslinked RNA immunoprecipitation (RIP-qPCR) experiment using *GFP-RZ-1C* and a *35S:GFP* control line. In order to test whether RZ-1C binds to nascent RNAs of *NCED6*, we performed RIP-qPCR using chromatin fraction (Figure 3c). The chromatin fraction was enriched by purifying nuclei, followed by stringent washes to remove non-chromatin-associated proteins (see Method) (Wu et al., 2016a; Zhu et al., 2020). The association between RZ-1C and nascent RNA was observed at *NCED6* and *ABI3* but not at a negative control gene *LEA*, suggesting *NCED6* and *ABI3* transcripts are bound by RZ-1C at the chromatin level.

### *rz-1 and swn clf* act synergistically in preventing seed germination and derepressing *NCEDs* expression

The above findings led to a hypothesis that RZ-1 functions in a silencing mechanism that operates at chromatin level. *NCEDs* are all intron-less, excluding the possibility that they are regulated through splicing. Among known gene silencing mechanisms, PRC2 mediated silencing is the most wide-spread at euchromatin in a range of different organisms. We noted that *NCEDs* remained in a repressed state post-germination (Figure 2b), which was accompanied by the presence of H3K27me3 based on public database (Zhang et al., 2007) (Supplementary Figure 3). We therefore examined whether *NCEDs* are also regulated by PRC2 during germination process. The PRC2-deficient mutant *swn clf* develops into callus post-germination and is therefore not viable; however, this distinct phenotype post-germination allows us to trace its seed germination speed. Consistent with a previous report (Bouyer et al., 2011), *swn clf* showed delayed germination similar to that observed for *rz-1b/1c* (Figure 4a). In addition, *NCEDs* are also derepressed to various extents in *swn clf* (Figure 4b).

**Figure 4.**
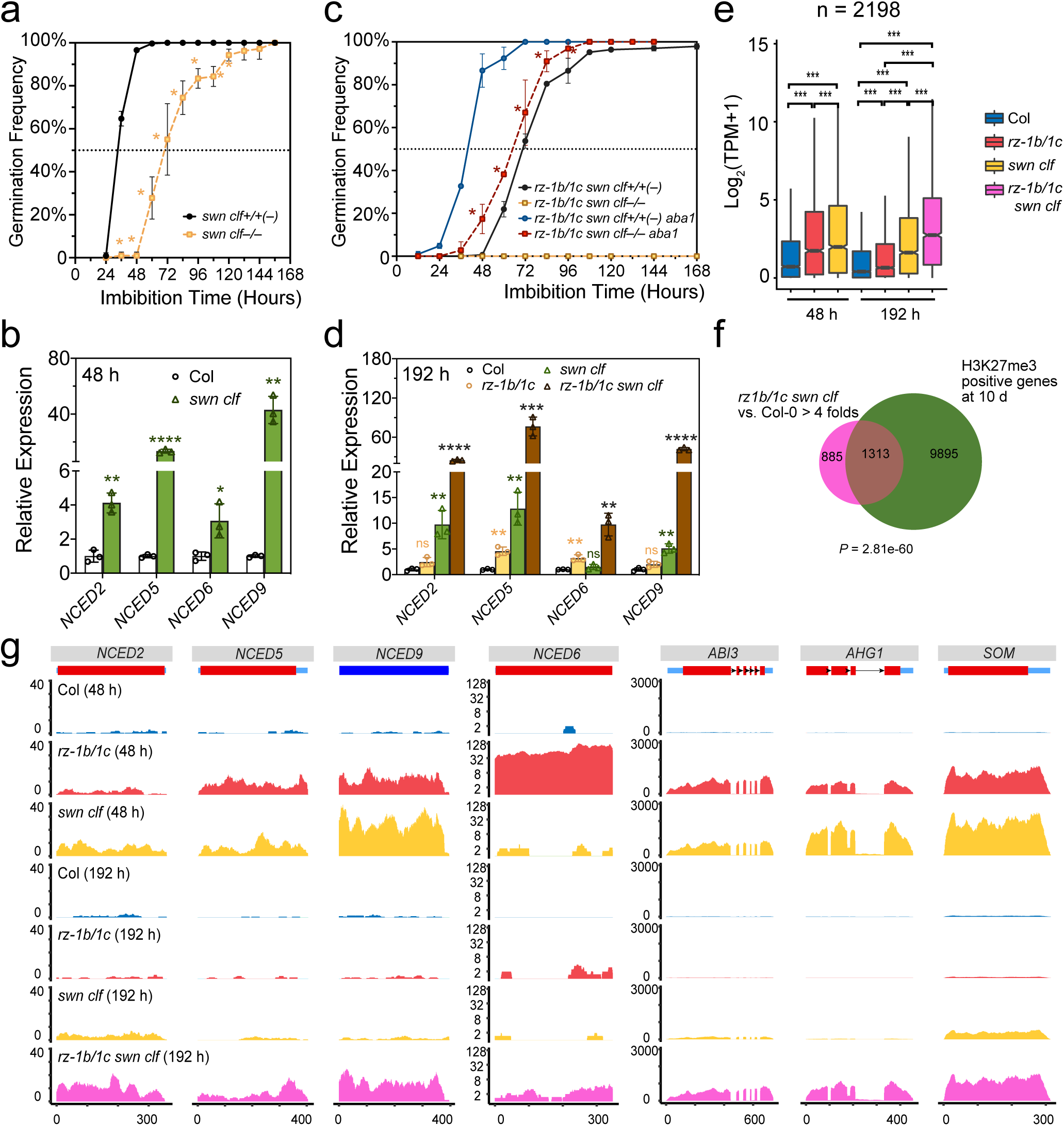
RZ-1C and PRC2 act synergistically to regulate seed germination at both the phenotypic and molecular levels. **a and c)** Seed germination frequency of *swn clf*, the *rz-1b/1c swn clf* quadruple mutant and the *rz-1b/1c swn clf aba1* quintuple mutant. Data are represented as mean ± SD from at least two biological replicates. * indicates significant difference between *swn clf+/+(–)* and *swn clf–/–* or between *rz-1b/1c swn clf–/– aba1* and *rz-1b/1c swn clf–/–* based on two-tailed Student’s *t*-test at *p* < 0.05. **b and d)** Gene expression analysis of *NCEDs* in different mutants after 48 h (**b**) or 192h (**d**) of imbibition. Fold changes over Col-0 are presented as mean ± SD from three biological replicates. *, **, *** and **** indicate statistical significance based on two-tailed Student’s *t*-test at *p*-values < 0.05, 0.01, 0.001 and 0.0001, respectively. ns = not significant. **e)** Box plot showing the distribution of expression levels of genes up-regulated by at least 4-fold in *rz-1b/1c swn clf* quadruple mutant compared to Col-0 at 192 h. TPM: transcript per million. *** indicate statistical significance based on Benjamini-Hochberg corrected *p*-value of pairwise t-test < 0.001. The box plot shows the median, the 25th and the 75th percentiles. **f)** Venn diagram showing the overlap between genes up-regulated in *rz-1b/1c swn clf* quadruple mutant and genes marked by H3K27me3 in Col-0 seedlings. **g)** Selected examples of synergy genes. Levels are mean of normalized sequencing depth (see Methods) from three biological replicates.

Next, we examined the genetic relationship between *rz-1b/1c* and *swn clf.* In an F2 population obtained from a cross between *swn clf* and *rz-1b rz-1c*, homozygote *rz-1b rz-1c swn clf* is missing from germinated individuals. However, surprisingly, individual genotyping of un-germinated seeds revealed that 97% of these seeds were *rz-1b rz-1c swn clf* homozygotes (Supplementary Figure 4). Therefore, the simultaneous loss of both RZ-1 and PRC2 blocks germination (Figure 4c). Importantly, this blocked germination depends on ABA biosynthesis, given that *rz-1b/1c swn clf aba1* quintuple mutant largely germinated normally (Figure 4c). In consistent, *NCEDs* were derepressed to much higher levels in *rz-1b/1c swn clf* than those in *rz-1b/1c* or *swn clf* double mutants after 8-day imbibition (Figure 4d). Therefore, RZ-1 and PRC2 synergize to repress *NCEDs* expression and to promote seed germination.

### RZ-1 and PRC2 synergize to repress lots of genes, including components of both ABA biosynthesis and signalling pathway

The above genetic data suggest that RZ-1 and PRC2 are neither in a linear genetic pathway nor in a completely parallel pathway. Instead, the data suggest both complexes may work cooperatively to achieve effective silencing of *NCEDs*. Next, we tested to what extent such a relationship between RZ-1 and PRC2 holds in a genome-wide scale. We profiled mRNA expression in Col-0, *rz-1b/1c* and *swn clf* at 48 h (seeds) and 192 h (seedlings) post-imbibition by mRNA-seq with biological triplicates (Supplementary Figure 5). The transcriptome of *rz-1b/1c swn clf* quadruple mutant was profiled only at 192 h as this mutant cannot be phenotypically separated from the segregating background plants at 48 h post-imbibition. The derepressed genes overlap significantly between *rz-1b/1c* and *swn clf* at both 48 h and 192h (Supplementary Figure 6), suggesting RZ-1 and PRC2 share common targets broadly. Next, to test the possible synergy between RZ-1 and PRC2, we compared the extent of gene derepression in *rz-1b/1c swn clf* to that in *rz-1b/1c* and *swn clf*. We identified 2198 genes that were upregulated more than 4 folds in *rz-1b/1c swn clf* comparing with Col-0 (Figure 4e). Remarkably, a synergistic effect between RZ-1 and PRC2 was observed in general for these genes (Figure 4e). Similar as we observed at *NCEDs*, the absence of either RZ-1 or PRC2 disrupts the effective silencing of these genes to various extent, while the absence of both complexes fully derepresses these genes and preventing their silencing over time (Figure 4e). Notably, 1313 out of 2198 genes are marked by H3K27me3 at seedling stage, indicating RZ-1 indeed synergize with PRC2 to repress many of the PRC2 targets (Figure 4f). Intriguingly, apart from *NCEDs*, various key ABA signalling genes that marked by H3K27me3 at seedling stage (public database (Zhang et al., 2007), also below paragraph) such as *ABI3*, *ABA-HYPERSENSITIVE GERMINATION 1* (*AHG1*) and *SOMNUS* (*SOM*) (Figure 4g) are also synergistically repressed by RZ-1 and PRC2, indicating both ABA biosynthesis and signalling pathway are coordinately repressed by the same set of mechanisms during germination. Taken together, the above showed RZ-1 and PRC2 synergize to repress lots of genes during seed germination.

### RZ-1 promotes silencing and the H3K27me3 at *NCED6* and other genes of which their H3K27me3 needs to be established through seed germination and early seedling development

The above data showed the presence of strong cooperativity between RZ-1 and PRC2 in gene silencing at transcriptome level. Next, we tested whether RZ-1 influences H3K27me3 levels at *NCED6*, *ABI3* and other targets shared by RZ-1 and PRC2. We profiled the H3K27me3 via ChIP-seq at 48 h (seeds) and 10 d (seedlings) post-imbibition in both Col-0 and *rz-1b/1c*. Good repeatability between biological replicates was observed (Supplementary Figure 7). In addition, the enrichment of H3K27me3 was observed at known PRC2 targets, such as *FLOWERING LOCUS* (*FT*), *SHOOT MERISTEMLESS* (*STM*) and *AGAMOUS* (*AG*), but was absent on the active gene *ACT7* (Figure 5a). Next, we looked at the H3K27me3 profiles at *NCED6*, *ABI3* and *AHG1* (Figure 5b). As expected, at these genes, H3K27me3 levels were significantly reduced in *rz-1b/1c* compared to Col-0 at 48 h (Figure 5b). At 10 days, the H3K27me3 is also reduced in *rz-1b/1c*, albeit to a less extent compared with 48 h (Figure 5b). In addition, H3K27me3 at these genes accumulated significantly from 48 h to 10 days (Figure 5b). Therefore, RZ-1 facilitates the accumulation of H3K27me3 at *NCED6*, *ABI3* and *AHG1* during germination and early seedling development.

**Figure 5.**
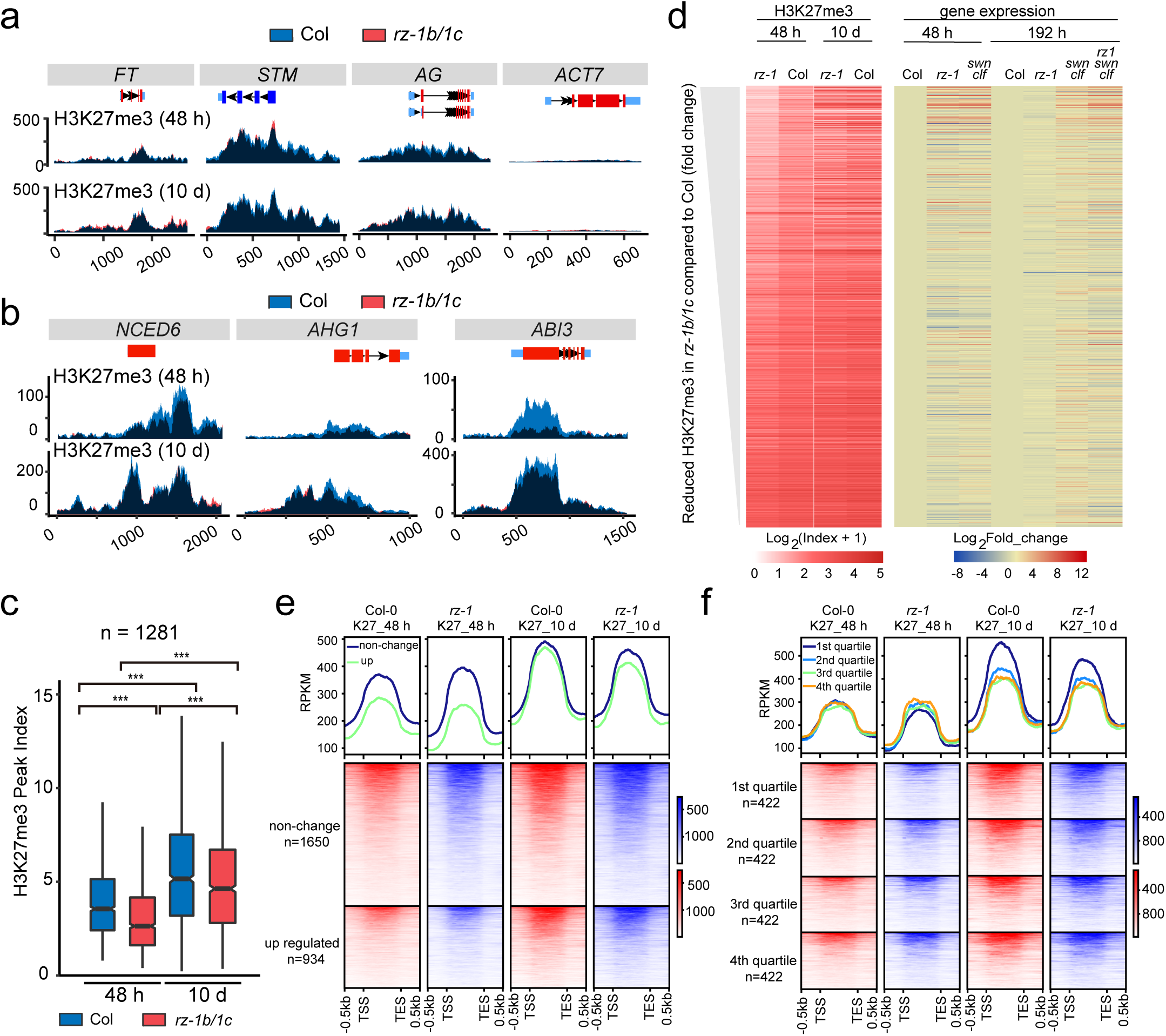
RZ-1 promotes silencing and H3K27me3 at genes whose H3K27me3 needs to be established through seed germination and early seedling development. **a and b)** H3K27me3 ChIP-seq profiles at individual control genes (a) or target genes (b) whose H3K27me3 levels are reduced in *rz-1b/1c* compared to Col-0 at 48 h post-imbibition. Blue indicates Col-0, red indicates *rz-1b/1c*, dark blue indicates the overlapping area between Col-0 and *rz-1b/1c*. Levels are mean of normalized sequencing depth (see Methods) from two (10 d) or three (48 h) biological replicates. **c)** Box plots of H3K27me3 levels of genes whose H3K27me3 levels are reduced in *rz-1b/1c* relative to Col-0 in 48-h imbibed seeds. *** indicate statistical significance based on Benjamini-Hochberg corrected *p*-value of pairwise t-test < 0.001. **d)** Patterns of H3K27me3 and expression levels of genes whose H3K27me3 levels are reduced in *rz-1b/1c* versus Col-0 at 48 h post-imbibition. Genes are ranked from top to bottom based on the extent of reduction in H3K27me3 levels in *rz-1b/1c*. Each line represents a single gene, and each track represents a specific histone mark at a certain time point (left panel) or gene expression level in each genotype (right panel). **e)** H3K27me3 profile of H3K27me3 marked genes whose mRNA levels are increased (> 2-fold) or unaltered (within ± 1.5 fold) in *rz-1b/1c* versus Col-0 at 48 h post-imbibition. **f)** H3K27me3 profile of H3K27me3 marked genes whose mRNA levels are increased (> 1.5 fold) in *rz-1b/1c* versus Col-0 at 48 h post-imbibition. Genes are equally divided into 4 quantiles according to their levels of derepression in *rz-1b/1c*, with the first quantile displays the highest level of derepression. For e) and f), genes are considered as H3K27me3 marked based on the presence of H3K27me3 peak on gene at 10 d seedling stage.

Next, we asked if RZ-1 is generally required for H3K27me3 accumulation during seed to seedling transition as we observed for *NCED6*, *ABI3* and *AHG1*. By differential peak calling, we identified 1281 genes with decreased H3K27me3 peak in *rz-1b/1c* compared with Col-0. For these genes, the H3K27me3 is decreased significantly in *rz-1b/1c* at both 48 h and 10 d, albeit the difference between two genotypes at 10 d is smaller compared with that at 48 h (Figure 5c). Notably, the absolute level of H3K27me3 increased at these genes from 48 h to 10 d (Figure 5c). In addition, sharp decreases of H3K27me3 in *rz-1b/1c* were frequently observed at genes with relatively low levels of H3K27me3 in Col-0 at 48 h (Figure 5d, towards the top of the heat map). Therefore, RZ-1 indeed facilitates a process of H3K27me3 accumulation that occurs at lots of genes during germination and early seedling development. Furthermore, we found PRC2 targets that were derepressed in *rz-1b/1c* generally display lower H3K27me3 compared with those not regulated by RZ-1 (Figure 5e), and the higher the levels of derepression in *rz-1b/1c*, the higher levels of H3K27me3 accumulation can be observed from 48 h to 10 d (Figure 5f). Taken together, the above data suggest RZ-1 is preferentially required for gene silencing at PRC2 targets when their H3K27me3 are not fully established, and the presence of RZ-1 also facilitates their H3K27me3 establishment in a timely manner.

### The interaction partners of RZ-1C at chromatin

To further explore the mechanism of RZ-1C mediated silencing, we looked into the protein interaction partners of RZ-1C in vivo at the chromatin level. In order to have an unbiased picture, we performed a crosslinked nuclear immunoprecipitation and mass spectrometry (CLNIP–MS) using imbibed seeds of *GFP-RZ-1C*. CLNIP is a reliable method for detecting both stable and transient interactions in vivo (Fang et al., 2019). Indeed, RZ-1C CLNIP captured a complex group of nuclear proteins with high confidence as partners of RZ-1C (Supplementary Figure 8, Supplementary Dataset 1). For example, 18 out of the 19 SR proteins, some of which are known direct interactors of RZ-1C (Wu et al., 2016b), were captured in this assay (Supplementary Figure 8, Supplementary Dataset 1). Apart from proteins involved in RNA metabolism (such as spliceosome components and m^6^A methyltransferase complex proteins), a few chromatin-related proteins were detected. Among these, we identified proteins that closely associate with PRC2, such as MSI1 (Derkacheva and Hennig, 2014; Costa and Dean, 2019) and components of CUL4-DDB1 complex (EMB1579, MSI4, DDB1A, DCAF1) (Dumbliauskas et al., 2011; Pazhouhandeh et al., 2011; Zhang et al., 2020)(Supplementary Figure 8, Supplementary Dataset 1). In addition, we also detected a few AL proteins (Supplementary Figure 8, Supplementary Dataset 1), which function as H3K4me3 readers (Lee et al., 2009). Histone deacetylase HDA19 and its closely related apoptosis- and splicing-associated protein (ASAP) complex (Supplementary Figure 8, Supplementary Dataset 1, SAP18 and ACINUS) were also detected. Notably, both *al* and *hda* mutants were reported to have similar molecular phenotypes as *rz-1b/1c* (see below paragraph, (Tanaka et al., 2008; Molitor et al., 2014)). Besides, RNA Polymerase II subunits and Pol II elongation factors were also detected (Supplementary Figure 8, Supplementary Dataset 1). The interactions between RZ1C and MSI1, HDA19 and AL2 were further independently confirmed by co-immunoprecipitation in *Arabidopsis* imbibed seeds in which MSI1, HDA19 and AL2 are fused with FLAG-tag and expressed under their native promoters respectively (Figure 6a, Supplementary Figure 9). Taken together, these results indicate that RZ-1 associates with multiple proteins including PRC2 peripheral proteins, H3K4me3 readers, histone deacetylases and the transcription apparatus during seed germination.

**Figure 6.**
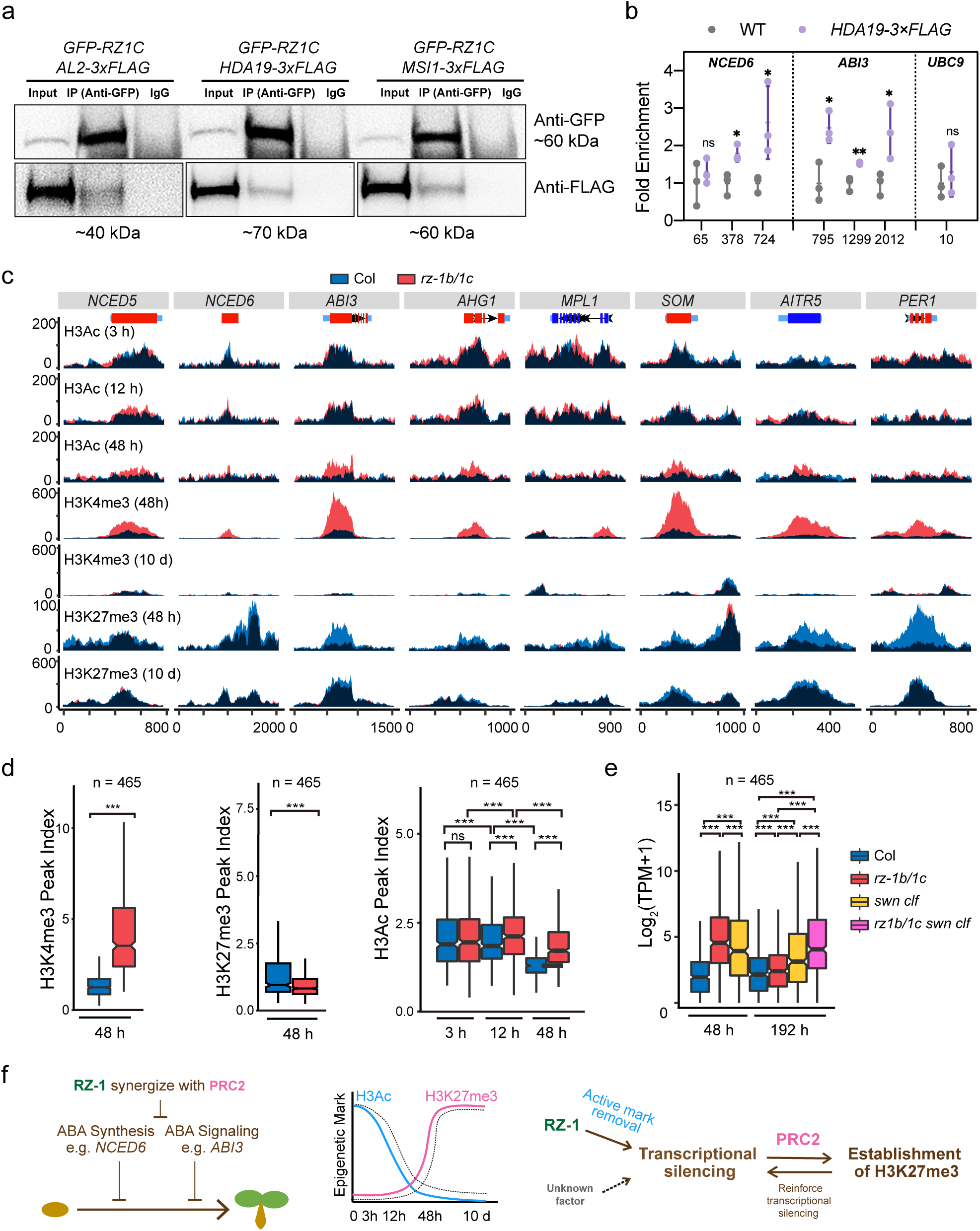
RZ-1 facilitates the removal of active histone marks and couples this process with PRC2 silencing. **a)** In vivo co-IP results validate interactions between RZ-1C and AL2, HDA19 or MSI1 in 24-h imbibed seeds. Nuclear extracts were used for IP with anti-GFP antibody or IgG (negative control). Western blots were performed using anti-GFP or anti-FLAG to show presence of both RZ-1C and target proteins. **b)** ChIP-qPCR results showing specific binding of HDA19 to *NCED6* and *ABI3* at chromatin level. *UBC9* serves as a negative control. qPCR data were first normalized to 1% input; fold enrichment was calculated by further normalization to WT control. Data were presented as mean ± SD from three biological replicates. *, ** and *** indicate statistical significance based on two-tailed Student’s t-test at p-values < 0.05, 0.01 or 0.001, respectively. ns = not significant. **c)** H3Ac, H3K4me3 and H3K27me3 profiles at selected genes that require RZ-1 for their H3 deacetylation and H3K4me3 suppression during germination. Blue, Col-0; red, *rz-1b/1c*; dark blue, overlapping area between Col-0 and *rz-1b/1c*. Levels are mean of normalized sequencing depth from two or three biological replicates. **d-e)** Box plots of the levels of histone modifications and expression of the top 25% genes with significantly higher H3K4me3 levels in *rz-1b/1c* compared to Col-0 at 48 h post-imbibition. The levels of H3K4me3, H3K27me3 and H3Ac are plotted in d) and the corresponding gene expression levels in different genotypes are plotted in e). *** indicate statistical significance based on Benjamini-Hochberg corrected *p*-value of pairwise t-test < 0.001. ns = not significant. **f)** A proposed working model of RZ-1 in promoting seed germination. Left, RZ-1 and PRC2 synergize to progressively silence ABA biosynthesis and signalling genes, as a result, seed germination occurs. Middle, Dynamics of histone modification accompanied with gene silencing during seed germination and early seedling growth, as exemplified at *NCED6*, *ABI3, AHG1* and others. H3Ac level gradually decreases while H3K27me3 level gradually increases. The dashed line indicates situation in *rz-1b/1c*. Right, A brief summary of the relationships between RZ-1 and PRC2. In part by promoting H3 deacetylation and suppression of H3K4me3, RZ-1 confers transcriptional silencing (e.g., *NCED6*, *ABI3* and *AHG1*) early during seed germination at PRC2 targets when their H3K27me3 are not yet established. This transcriptional silence state likely facilitates the H3K27me3 modification mediated by PRC2. The establishment of H3K27me3 further reinforces the transcriptional silencing to achieve an epigenetically stable state. In the absence of RZ-1, unknown factor likely presents to transcriptionally off switch the locus and enables H3K27me3 establishment, albeit at a slower rate.

### RZ-1 promotes transcriptional silencing of *NCED6* and other genes in part by promoting histone deacetylation and suppression of H3K4me3

The data showed in Figure 3 to Figure 5 suggests RZ-1 confers silencing of *NCED6* and other genes when the H3K27me3 has not been fully established yet. Given RZ-1 associates with HDA19 and ALs, we then tested if RZ-1 confers transcriptional silencing early during seed germination at *NCED6* and other genes involving histone deacetylation and suppression of H3K4me3. We found *hda19* single mutant did not show defective germination despite the upregulation of a few ABA genes (*NCED5*, *NCED9* and *ABI5*), likely due to functional redundancy or genetic compensation from other histone deacetylases (Supplementary Figure 10a, b). Indeed, inhibition of histone deacetylases by trichostatin A (TSA) treatment did lead to defective germination and derepression of *NCED2, NCED5, NCED6*, *NCED9* and *ABI5* in addition to *ABI3* (Tanaka et al., 2008) during imbibition (Supplementary Figure 10c, d), indicating a positive role of histone deacetylation in silencing these genes. In consistent, using ChIP-qPCR, we found HDA19 associates with *ABI3* and *NCED6* chromatin, but not with *UBC9*, in imbibed seeds (Figure 6b). For H3K4me3 reader ALs, we found *al3* mutant showed delayed germination and derepression of *NCEDs* (Supplementary Figure 10a, b), suggesting ALs are likely involved in the suppression of ABA synthesis, in addition to their known function in suppressing *ABI3* (Molitor et al., 2014). We further profiled H3K4me3 and H3Ac in Col-0 and *rz-1b/1c* through ChIP-seq (Supplementary Figure 7). In order to inspect potential deacetylation of histone H3, we profiled H3Ac at 3 h, 12 h and 48 h post imbibition. As expected, at *NCED6*, *AHG1, ABI3, SOM1* and other individual genes, H3K4me3 and H3Ac levels are significantly higher in *rz-1b/1c* compared with Col-0 at 48 h (Figure 6c).

Importantly, at these genes, H3Ac levels are decreased from 3 h to 48 h and such a process is delayed in *rz-1b/1c* (Figure 6c). Therefore, RZ-1 confers transcriptional silencing of *NCED6* and other genes early during seed germination at least in part through promoting the H3 acetylation and suppression of H3K4me3.

Finally, we examined the link between increased H3Ac/H3K4me3 and altered H3K27me3/gene expression in *rz-1b/1c*. We identified genes with significantly increased H3Ac or H3K4me3 levels in *rz-1b/1c* at 48 h post-imbibition and profiled all of the histone modification and expression data accordingly. Both groups of genes showed a significant decrease in H3K27me3 levels in *rz-1b/1c* (Figure 6d and Supplementary Figure 11). In addition, both groups of genes showed a clear RZ-1-enhanced deacetylation pattern during germination process (Figure 6d and Supplementary Figure 11). Furthermore, both groups of genes showed sharply upregulated expression not only in *rz-1b/1c* but also in *swn clf* at 48 h post-imbibition, and that synergy between *rz-1b/1c* and *swn clf* was observed in the 192 h dataset (Figure 6e and Supplementary Figure 11). Taken together, these findings indicate that RZ-1 promotes histone deacetylation and suppression of H3K4m3 during seed germination. These series of activities confer a transcriptional silence state, such a state in one hand compensates for PRC2 when H3K27me3 is not fully established, and on the other hand facilitates the timely establishment of H3K27me3. The collective efforts of RZ-1 and PRC2 at both ABA biosynthesis and signalling genes then ensure germination (Figure 6f).

## Discussion

In this study, we reported a gene silencing mechanism that is critical for seed germination in *Arabidopsis*. RZ-1 promotes the transcriptional silencing of key ABA biosynthesis and signalling genes (Figure 6f), involving its interaction with nascent RNAs, histone deacetylation and suppression of H3K4me3, largely during seed germination (Figure 6f). RZ-1 compensates for PRC2 and also cooperates with PRC2 such that RZ-1 mainly exerts its silencing function when H3K27me3 is not fully established. In addition, RZ-1 enhances the timely establishment of H3K27me3, which likely initiates during germination and finishes during early seedling development, when the transcription of the locus is switched off (Figure 6f). We propose RZ-1 may facilitate PRC2 action mainly through promoting active mark removal given transcription was proposed as an opposing state of PRC2 silencing (Berry et al., 2017). Indeed, the presence of active chromatin marks such as H3K4me3 inhibits PRC2 activity (Schmitges et al., 2011). The transcriptionally repressed state mediated by RZ-1 then facilitates the H3K27me3 establishment likely indirectly (Figure 6f). The establishment of H3K27me3 at the locus further feeds back to reinforce the transcriptional silencing and confers an epigenetically stable state. Notably, a mechanism that RZ-1 works through recruiting PRC2 is unlikely given RZ-1 mainly associates with PRC2 peripheral MSI1 and CUL4-DDB1 complex but not PRC2 catalytic core. The relationship between RZ-1 and PRC2 are best demonstrated at *NCED6* and *ABI3*, both of which contribute to the delayed germination phenotype in *rz-1b/1c*. In the absence of PRC2, *ABI3* and *NCED6* are not fully derepressed (Figure 4g). This is due to RZ-1, the presence of which is sufficient to confer a transcriptionally repressed, likely metastable state, when the H3K27me3 are not established. On the other hand, in the absence of RZ-1, other unknown protein likely functions to transcriptionally switch off the locus and allows the establishment of H3K27me3, albeit at a slower rate. Overall, RZ-1 and PRC2 are in the same genetic pathway such that they tightly cooperate, while their exact working mechanism at the molecular level is different.

Our genetic data show that the silencing of *NCED6*, in particular, contributes most to timely germination in an RZ-1 dependent manner (Figure 3a, b). Current evidence suggests that RZ-1 directly represses *NCED6*, as RZ-1C targets to the nascent RNA of *NCED6* (Figure 3c). In addition, the expression of known regulators of *NCED6* is either unchanged or altered in a way supposed to repress *NCED6* in *rz-1b/1c* (Supplementary Figure 12). In addition to *NCED6*, the silencing mechanism we discovered also operates for several key ABA signalling genes such as *ABI3*, *AHG1* and *SOM1* (Figure 4g, 6c). Therefore, in non-dormant seeds, the ABA pathway is robustly and coordinately silenced at multiple key nodes during imbibition, thereby ensuring seed germination (Figure 6f).

For non-dormant seeds, it has been known for decades that ABA level drops dramatically within the first few hours of imbibition, which is attributed to rapid ABA catabolism (Kushiro et al., 2004; Okamoto et al., 2006; Liu et al., 2009). The finding that *NCEDs* are not in a pre-silenced state before germination was unexpected; instead, they are silenced progressively throughout the germination process, which likely also contributes to the decline in ABA content along the germination process. Such progressive silencing might be beneficial for the flexibility of seed germination. Indeed, unfavourable conditions during imbibition could trigger secondary dormancy to prevent germination, a process that is also regulated by ABA (Ibarra et al., 2016).

We demonstrated that *NCED6* and many other genes are transcriptionally silenced through progressive histone deacetylation during seed germination (Figure 4g, 6c to 6e), followed by PRC2-mediated establishment of H3K27me3. Interestingly, histone deacetylation is the first histone modification that is regulated during X-chromosome inactivation (Zylicz et al., 2019). In plant, histone deacetylation is important for the transcriptional off-switch of *Arabidopsis FLC* gene through vernalization (Questa et al., 2016). Therefore, histone deacetylation might be functionally conserved and be a common prior to facultative heterochromatin formation in a range of different systems.

The exact contribution of RNA-binding in RZ-1 mediated silencing remains to be further investigated in the future. Current data indicate RZ-1 likely functions in multiple aspects of RNA metabolism at the chromatin level. RZ-1C associates with nascent RNA of *NCED6* and *ABI3* (Figure 3c). Notably, apart from HDA19, H3K4me3 readers and PRC2 related proteins, the majority of RZ-1-associated proteins are RNA metabolism-related proteins (Supplementary Figure 8, Supplementary Dataset 1). Additionally, Pol II subunit, Pol II elongation factors and m^6^A writer complex were also detected (Supplementary Figure 8, Supplementary Dataset 1), which surprisingly resembles the case of SPEN protein during X-chromosome inactivation (Dossin et al., 2020). How exactly RZ-1 determines its primary targets such as *NCED6* is also unknown. The sequence specific binding of RNA by RZ-1 is unlikely to be sufficient for its recruitment, given that AAAGA (Wu et al., 2016b; Zhu et al., 2020) or purine rich motif is wide-spread within the genome. In addition to histone deacetylation, it is likely that RZ-1 also integrates multiple coordinated activities such as RNA metabolism at the locus and also transcriptional initiation and elongation; these series of activities shut down the locus and further cooperate with PRC2 to stably silence the locus. Notably, *NCED6* is intronless and there is no sign of defective splicing at *ABI3* in *rz-1b/1c* judged by mRNA-seq data, suggesting splicing is unlikely the key for RZ-1 mediated silencing. Targeted genetic screening would be required in future to fully dissect this mechanism. In conclusion, we uncovered the progressive chromatin silencing of ABA biosynthesis and signalling genes mediated cooperatively by RZ-1 and PRC2, such a mechanism gates seed germination in *Arabidopsis*.

## Methods

### Plant Materials and Growth Conditions

The *Arabidopsis thaliana* materials used in this study were as follows: the Col-0 ecotype was used as the WT control; the *rz-1b/1c* double mutant and *GFP-RZ-1C* complemented line were described previously(Wu et al., 2016b). The *RZ-1C: GUS-GFP-RZ-1C* construct was generated by modifying the pEASY-GFP-RZ-1C construct used to create the *GFP-RZ-1C*-complemented line as described previously(Wu et al., 2016b). In brief, the *GUS* fragment was amplified and joined together with *GFP-RZ-1C* in a linearized vector using Gibson Assembly. The resulting *RZ-1C: GUS-GFP-RZ-1C* vector was subcloned into binary vector pH7FWG0 using Gibson Assembly. For generation of *AL2-3×FLAG*, *HDA19-3×FLAG* and *MSI-3×FLAG* constructs, their genomic sequences from 2kb upstream of transcription start site to where immediately before stop codon were amplified by PCR, and inserted into pk2GW7 immediately before the sequence coding 3xFLAG, using Gibson Assembly. For all transgenic lines, the respective final construct was first transformed into *Agrobacterium tumefaciens* GV3101 and then transformed into *rz-1b/1c* plant (for RZ-1 related constructs) or *GFP-RZ-1C rz-1b/1c* plant (for FLAG-tagged line) using the floral dip method. Successful transformants were identified by antibiotic selection and PCR confirmation. ABA-related mutants were purchased from Nottingham Arabidopsis Stock Centre (NASC), including *aba1-6*, *nced2* (SALK_026541C), *nced5* (GK_380E07), *nced6* (WiscDsLox388C03), *nced9* (GK-624A05) and *abi3* (SALK_003216). *hda19(SALK_027241C)* and *al3(SALK_080056C)* mutants were purchased from Arashare. *swn-7 clf-28/CLF* was kindly provided by Dr. Caroline Dean. The triple, quadruple and quintuple mutants used in this study were generated by genetic crossing of the above mutants and bulking in the laboratory. The primers used for genotyping are presented in Supplementary Dataset 2.

In most experiments, seeds were sterilized in 15% (v/v) NaClO for 10 min and sown on half-strength Murashige and Skoog (MS) medium with 1% (w/v) agar. For hormone/inhibitor treatments, seeds were sown on half-strength MS medium supplemented with the proper doses of chemicals. For germination analysis, seeds were sown on plates and transferred immediately to a growth chamber without stratification (16-hour light/8-hour dark cycle, 22°C). The germination status (radical emergence) of each seed was determined at the designated time points under a stereomicroscope.

### ABA quantification

Three biological replicates of seed samples were used for this analysis. Approximately 30 mg dry seeds or 150 mg imbibed seeds per sample were collected over a time course (0, 3, 6, 12, 24, 36 and 48 h post-imbibition), flash-frozen and ground into a fine powder in liquid nitrogen. The samples were freeze-dried in a Labconco FreeZone freeze dryer, and the dry weight of each sample was recorded for normalization. ABA quantification was performed at Kunming Institute of Botany by HPLC-MS/MS as described in (Wu et al., 2007; Qi et al., 2016) with minor modifications. Briefly, 1 ml ethyl acetate spiked with 10 ng D6-ABA as an internal standard was added to each sample. The sample was immediately vortexed for 10 min and centrifuged at maximum speed for 10 min at 4°C. After transferring the supernatant into a fresh tube, the pellet was re-extracted with 0.5 ml of ethyl acetate, and the supernatants were combined after centrifugation. The sample was dried in a vacuum concentrator and the residue resuspended in 0.5 ml 70% methanol (v/v). After centrifuging the sample to clarify the phases, the supernatant was carefully transferred into a glass vial for HPLC-MS/MS analysis (LCMS-8040, Shimadzu). Measurements were conducted using a LC-20AD liquid chromatography system (Shimadzu). 10 μl of each sample was injected onto an ODS column (Shim-pack XR-ODS III, 1.6 μm, 75 × 2 mm) at a flow rate of 0.3 ml/min. The ABA level in each sample was determined by comparing its peak area with that of the internal standard. Final ABA concentration was calculated by normalizing to the dry weight of each sample before extraction.

### Histochemical GUS staining assay

*GUS-GFP-RZ-1C* was constructed as described in the Plant Materials section. Seeds were sown on half-strength MS medium for 24 or 48 h. Imbibed seeds were vacuum infiltrated with X-Glc staining solution [1 mg X-Glc in 1 ml GUS buffer: 0.5 M Na_2_HPO_4_, 0.5 M NaH_2_PO_4_, 100 mM K_3_Fe(CN)_6_ and 100 mM K_4_Fe(CN)_6_] for 10 minutes, followed by incubation at 37°C overnight. The seeds were briefly de-stained in 70% (v/v) ethanol and observed and photographed under a stereomicroscope (Zeiss Stemi 508).

### Total RNA extraction and quantitative RT-PCR

Approximately 30 mg dry seeds or 100 mg imbibed seeds were collected, flash-frozen and finely ground in liquid nitrogen for total RNA extraction using a HiPure Plant RNA Mini Kit following the manufacturer’s protocol (Magen). DNA contamination was removed with a RapidOut DNA Removal Kit (Thermo Scientific). For each sample, 1 µg total RNA was used to synthesize cDNA using M-MLV Reverse Transcriptase (Promega) and Oligo d(T)_18_. A 50-fold dilution of cDNA was used for qPCR using a real-time thermal cycler (qTOWER³ 84, Analytik Jena) and SYBR Green qPCR Master Mix (Roche). *UBC9* was used as an internal reference control. Values in the figures are mean ± SD from at least three biological replicates and are displayed as per *UBC9* level or fold change relative to Col-0. The primer sequences are presented in Supplementary Dataset 2.

### mRNA purification and sequencing library construction

mRNA-seq was performed as described previously(Zhu et al., 2020). 5 µg total RNA was used for mRNA purification using VAHTS mRNA capture beads following the manufacturer’s protocol (Vazyme). 50 ng mRNA was used to construct a sequencing library using the dUTP method and with a NEBNext^®^ Ultra II Directional RNA Library Prep Kit for Illumina following the manufacturer’s protocol. Briefly, mRNA was fragmented and random primed for first-strand cDNA synthesis in the presence of dUTP. Second-strand cDNA was synthesized, and double-stranded cDNA was ligated to adaptors. Adaptor-ligated dsDNA was treated to remove the first strand of DNA to maintain the strand information. The library was then amplified by PCR with Universal PCR Primer/i5 Primer and Index (X) Primer /i7 Primer. The resulting library was purified, quantified and used for pair-end Illumina (PE150) sequencing. mRNA-seq at 24 h was performed once and only used for the initial screening of ABA-related genes with altered expression in *rz-1b/1c* (Figure 2a); the results were validated by qPCR (Figure 2c, d). All of the mRNA-seq data presented in Figure 4 to 6 were obtained using three biological replicates.

### RNA Immunoprecipitation (RIP)-qPCR

RIP-qPCR was performed as described previously (Zhu et al., 2020) with some modifications. 12-h imbibed seeds of the GFP-RZ-1C and 35S-GFP transgenic lines were crosslinked under a vacuum with 1% formaldehyde in 1 × PBS for 15 min. Glycine was added to quench the extra formaldehyde. The samples were washed several times in distilled water, flash-frozen in liquid nitrogen and ground into a fine powder. For RIP of the chromatin fraction, nuclei were enriched using Honda buffer [20 mM HEPES, 0.44 M sucrose, 1.25% (w/v) Ficoll, 2.5% (w/v) Dextran T40, 10 mM MgCl_2_, 0.5% (v/v) Triton X-100, 2 mM DTT, 1 × protease inhibitor cocktail, 100 ng/μl tRNA, 20 U/ml RNase Inhibitor] as described previously(Zhu et al., 2020). The pellet obtained from each 1.2 g plant sample was resuspended in nuclei resuspension buffer [25 mM Tris-HCl pH 7.5, 100 mM NaCl, 0.5 mM EDTA, 50% (v/v) glycerol, 2 mM DTT, 2× protease inhibitor cocktail, 200 ng/μl tRNA, 40 U/ml RNase inhibitor] following two sequential washes with urea washing buffer [25 mM Tris-HCl pH 7.5, 300 mM NaCl, 0.5 mM EDTA, 1 M urea, 1% (v/v) Tween 20, 2 mM DTT, 2 × protease inhibitor cocktail, 200 ng/μl tRNA, 40 U/ml RNase inhibitor]. The resulting chromatin pellet was washed once in nuclei resuspension buffer to remove traces of urea, pelleted and resuspended in nuclear lysis buffer. The chromatin samples were sonicated, centrifuged and the supernatant subjected to immunoprecipitation. The chromatin fractions were diluted in IP dilution buffer [50 mM Tris-HCl pH 8.0, 150 mM NaCl, 1 mM EDTA, 1% (v/v) Triton X-100] for immunoprecipitation. 1/100 diluted samples were saved as input. 2.5 ml diluted samples were incubated with 5 μg anti-GFP antibody (Abcam, ab290) and 50 μl Protein A Agarose/Salmon Sperm DNA (Millipore) at 4°C for 2 h. After IP, the beads were washed four times in IP dilution buffer and once in TE buffer. The samples on beads, together with the previously saved inputs, were reverse crosslinked overnight in elution buffer [50 mM Tris-HCl pH 8.0, 2 mM EDTA, 0.2% (v/v) SDS, 200 ng/μl tRNA and 20 mg/ml Proteinase K]. RNA was recovered from the samples using the TRIzol method, treated with DNase and precipitated for reverse transcription using target-gene-specific reverse primer mix. The synthesized cDNA was diluted 50-fold for qPCR analysis. Values from IP samples were normalized to the respective input and displayed as mean ± SD from four biological replicates. *35S-GFP* was used as the background control. The primer sequences are presented in Supplementary Dataset 2.

### ChIP-seq library preparation

Histone ChIP-seq was performed as described previously with some modifications(Wu et al., 2016a; Hagai et al., 2018). In brief, nuclei were extracted from 0.5 g samples using Honda buffer. The resulting pellet was resuspended in nuclear lysis buffer [50 mM Tris-HCl pH 7.5, 10 mM EDTA pH 8.0, 2 × protease inhibitor cocktail and 0.5% (v/v) SDS], sonicated, centrifuged, diluted and subject to immunoprecipitation using Dynabeads protein A (Invitrogen) pre-bound with the following antibodies: anti-trimethyl-Histone H3 (Lys27) antibody (Millipore: 07-449), Histone H3K4me3 antibody (Active Motif: 39915) and anti-acetyl-Histone H3 antibody (Millipore: 06-599). Following IP, the beads were washed sequentially in washing buffer A [20 mM Tris-HCl pH 7.5, 2 mM EDTA pH 8.0, 150 mM NaCl, 1% (v/v) Triton X-100], washing buffer B [20 mM Tris-HCl pH 7.5, 2 mM EDTA pH 8.0, 500 mM NaCl, 1% (v/v) Triton X-100], washing buffer C [10 mM Tris-HCl pH 8.0, 1 mM EDTA pH 8.0, 250 mM LiCl, 1% (v/v) NP40, 0.5% (w/v) Na-deoxycholate], washing buffer D [10 mM Tris-HCl pH 7.5, 1 mM EDTA pH 8.0, 1% (v/v) Triton X-100] and 1xTE buffer (10 mM Tris-HCl pH 8.0, 1 mM EDTA pH 8.0, 50 mM NaCl).

The resulting samples were used for on-bead adaptor ligation using Tn5 Transposase (Vazyme). The samples were reverse crosslinked in elution buffer [50 mM Tris-HCl pH 8.0, 10 mM EDTA pH 8.0 and 1% (v/v) SDS] at 65°C overnight. DNA was recovered using a MinElute PCR Purification kit (Qiagen). The sequencing library was amplified by PCR using 2 × NEBNext High-Fidelity PCR Master Mix and N5xx/N7xx primers from a Nextera Index Kit. The resulting library was purified, quantified and used for pair-end Illumina (PE150) sequencing.

### ChIP-qPCR

ChIP-qPCR was performed as described previously with some modifications (Wu et al., 2016a). Briefly, chromatin was prepared from 1% formaldehyde crosslinked seeds by using Honda buffer. Following sonication and dilution of the supernatant, the samples were immunoprecipitated using anti-FLAG^®^ M2 magnetic beads (Millipore: M8823) at 4°C overnight for HDA19 ChIP experiment. The beads were washed and incubated in elution buffer at 65°C overnight and treated with proteinase K for 2 h at 55°C for reverse crosslinking. Immunoprecipitated DNA was recovered by phenol: chloroform extraction followed by isopropanol precipitation. qRT-PCR data were first normalized to respective input and then to corresponding control samples. The primer sequences used for ChIP-qPCR are presented in Supplementary Dataset 2.

### Crosslinked nuclear immunoprecipitation and mass spectrometry

CLNIP–MS was performed as described previously (Fang et al., 2020). In brief, nuclei were extracted from 24-h imbibed GFP-RZ-1C seeds that had been crosslinked with 1% formaldehyde as described above. Each 2 g sample was combined with 8 ml lysis buffer [20 mM Tris-HCl pH 7.5, 20 mM KCl, 2 mM EDTA pH 8.0, 2.5 mM MgCl_2_, 25% (v/v) glycerol and 250 mM sucrose], homogenized and filtered through one layer of Miracloth. After centrifuging the sample and washing in NRBT buffer [20 mM Tris-HCl pH 7.5, 2.5 mM MgCl_2_, 25% (v/v) glycerol and 0.2% (v/v) Triton X-100] three times, the pellet was resuspended in RIPA buffer [1 × PBS, 1% (v/v) NP-40, 0.5% (w/v) sodium deoxycholate and 0.1% (v/v) SDS] for subsequent sonication. The samples were centrifuged and the supernatants used for immunoprecipitation with GFP antibody (Abcam: ab290) and Dynabeads Protein A (Invitrogen). Immunoprecipitates on beads were washed sequentially in low salt [20 mM Tris-HCl pH 8.0, 2 mM EDTA pH 8.0, 150 mM NaCl, 0.1% (v/v) SDS, 1% (v/v) Triton X-100,], high salt (same recipe as low salt buffer except with 500 mM NaCl) and TE buffer (10 mM Tris-HCl pH 8.0, 1 mM EDTA pH 8.0) and boiled in 1 × NuPAGE LDS sample buffer for 30 min at 95°C. The samples were separated on a NuPAGE denaturing gel (10%, Bis-Tris, 1.0 mm), and regions excluding heavy and light chains were recovered and subjected to in-gel trypsin digestion as described previously(Shevchenko et al., 2006).

Following digestion, the supernatants were dried in a SpeedVac concentrator (Thermo Fisher) and analysed by LC-MS using Nano LC-MS/MS (Dionex Ultimate 3000 RLSCnano System) interfaced with Q Exactive HF. Mass spectrometric raw files were analysed using the peptide search engine Andromeda (Cox et al., 2011) integrated into the MaxQuant environment(Cox and Mann, 2008). The resulting MS/MS spectra were searched against the TAIR10 protein database. LFQ (label free quantitation), iBAQ (intensity Based Absolute Quantification) and unique peptides were considered for quantification. The reported values for protein quantification were processed using statistical methods. Briefly, GFP-RZ-1C LFQ values were compared to those of the wild-type control by Student’s *t*-test, as shown in Supplementary Figure 8. Significantly enriched proteins bound by RZ-1C were further analysed and categorized manually. The selected RZ-1C-associated proteins with LFQ ratios > 2 are presented in Supplementary Dataset 1.

### mRNA-Seq data analysis

A summary of mRNA-seq data is presented in Supplementary Dataset 3. The raw data were cleaned using the Trimmomatic(Bolger et al., 2014) package (version 0.39). The adapters, Ns and low-quality bases were removed, and any reads that were smaller than 36 bp after adaptor removal were also excluded. The clean reads were mapped to the TAIR10 genome using HISAT2 (Kim et al., 2019) (version 2.2.0) with default parameters. The uniquely mapped reads were retained for further processing using SAMtools (Li et al., 2009) (version 1.3.1). The read numbers were counted using featureCounts (Liao et al., 2014) (version 2.0.0), and transcripts per million (TPM) values were calculated using TPMCalculator (Vera Alvarez et al., 2019) (version 0.0.3). Differentially expressed genes were identified by DESeq2 (Love et al., 2014) (version 1.28.1), with a fold change value no less than 2 and a corrected *p* value no more than 0.05. The statistical test of the overlap between two gene groups was performed using the GeneOverlap package (http://shenlab-sinai.github.io/shenlab-sinai/). The box plots indicate the first and third quartiles (boxes), the median (horizontal bands) and the 95% confidence interval of the median (notches). The upper whisker extends from the hinge to the largest value no further than 1.5 * IQR from the hinge (where IQR is the inter-quartile range, or distance between the first and third quartiles). The lower whisker extends from the hinge to the smallest value at most 1.5 * IQR of the hinge. To display mRNA-seq data at the single gene levels, sequencing depths were calculated using a 5 bp non-overlapping sliding window and normalized to the total sequencing depth; the average value from three biological replicates is shown.

### ChIP-Seq data analysis

A summary of the ChIP-seq data is presented in Supplementary Dataset 3. The initial quality check and filtering of the ChIP-seq data were performed as described for mRNA-seq. The clean reads were aligned to the TAIR10 genome using Bowtie 2 (Langmead and Salzberg, 2012) (version 2.3.5.1) with default parameters. The reads with mapping quality < 20 were removed using SAMtools (Li et al., 2009) (version 1.3.1). PePr (Zhang et al., 2014) (version 1.1.24) was used for peak calling against the input considering variations between biological replicates and with a window size of 100 bp. Differential peaks between samples were identified using the “diff” function of PePr. The identified peaks were filtered according to the following criteria: (i) peak length *≥* 400 bp; (ii) fold change value (over input or between samples) *≥* 1.5; and (iii) *p* value no more than 1e-5. Genes that overlapped with existing peaks were annotated as either genes overlapping > 200 bp with existing peaks or the overlapping region between a peak and a gene accounted for more than 50% of the peak or the gene. The index value of a peak or gene was calculated as Index = Depth _peak_200_ / Depth _average_, where the Depth _peak_200_ is the average sequencing depth of the 200 bp region around the top of a peak, and the Depth _average_ is the average sequencing depth of the whole genome. For metagene profiling, RPKM was calculated with a 10 bp sliding window, and visualization was performed using plotProfile in deepTools. Analysis of the chromosome distribution of altered H3K27me3 peaks was performed using the RIdeogram package (https://cran.r-project.org/web/packages/RIdeogram/). The box plots for ChIP-seq data were defined the same as in mRNA-seq data analysis section. To display ChIP-seq data at a single gene, sequencing depth was calculated using 5 bp non-overlapping sliding windows and normalized to the total sequencing depth; the average values from the available biological replicates are shown.

## Data availability

The mRNA-seq and ChIP-seq datasets generated in this study have been deposited into the SRA database at NCBI under BioProject IDs PRJNA656062 and PRJNA662156, respectively.

## Acknowledgments

We thank Dr. Jinfeng Qi and Dr. Jianqiang Wu (KIB, China) for their generous help on ABA content measurement. We thank Dr. Caroline Dean (JIC, UK), Dr. Hongwei Guo (SUSTech, China) and Dr. Yong Xiang (AGIS, China) for their critical comments and suggestions on this work. This work was supported by the National Natural Science Foundation of China (31771365 to Z.W., 31900248 to D.Y. and 31800268 to D.Z), Guangdong Innovation Research Team Fund (2016ZT06S172), the Shenzhen Sci-Tech Fund (KYTDPT20181011104005) and Key Laboratory of Molecular Design for Plant Cell Factory of Guangdong Higher Education Institutes (2019KSYS006).

## Author Contributions

Z.W., D.Y., D.Z., L.Q., X.C., J.D., Y.W., and M.C. conceived the study. D.Y. conducted the majority of the experiments. F.Z., conducted the majority of bioinformatic analysis. X.K. generated the *RZ-1C: GUS-GFP-RZ-1C* constructs and corresponding transgenic lines. D.Z. and X.C. developed ChIP-seq protocol. Z.W. and D.Y. wrote the paper with input from all authors.

---

**Supplementary Dataset 1. Summary of IP-MS results**

**Supplementary Dataset 2. Primers used in this study**

**Supplementary Dataset 3. Summary of sequencing data**

## References

Barros-Galvao, T., Dave, A., Gilday, A.D., Harvey, D., Vaistij, F.E., and Graham, I.A. (2020). ABA INSENSITIVE4 promotes rather than represses PHYA-dependent seed germination in Arabidopsis thaliana. New Phytol 226, 953–956.

Berry, S., and Dean, C. (2015). Environmental perception and epigenetic memory: mechanistic insight through FLC. Plant J 83, 133–148.

Berry, S., Dean, C., and Howard, M. (2017). Slow Chromatin Dynamics Allow Polycomb Target Genes to Filter Fluctuations in Transcription Factor Activity. Cell Syst 4, 445–457 e448.

Bolger, A.M., Lohse, M., and Usadel, B. (2014). Trimmomatic: a flexible trimmer for Illumina sequence data. Bioinformatics 30, 2114–2120.

Bouyer, D., Roudier, F., Heese, M., Andersen, E.D., Gey, D., Nowack, M.K., Goodrich, J., Renou, J.P., Grini, P.E., Colot, V., and Schnittger, A. (2011). Polycomb repressive complex 2 controls the embryo-to-seedling phase transition. PLoS Genet 7, e1002014.

Brockdorff, N., Bowness, J.S., and Wei, G. (2020). Progress toward understanding chromosome silencing by Xist RNA. Genes Dev 34, 733–744.

Costa, S., and Dean, C. (2019). Storing memories: the distinct phases of Polycomb-mediated silencing of Arabidopsis FLC. Biochem Soc Trans 47, 1187–1196.

Cox, J., and Mann, M. (2008). MaxQuant enables high peptide identification rates, individualized p.p.b.-range mass accuracies and proteome-wide protein quantification. Nat Biotechnol 26, 1367–1372.

Cox, J., Neuhauser, N., Michalski, A., Scheltema, R.A., Olsen, J.V., and Mann, M. (2011). Andromeda: a peptide search engine integrated into the MaxQuant environment. J Proteome Res 10, 1794–1805.

Derkacheva, M., and Hennig, L. (2014). Variations on a theme: Polycomb group proteins in plants. J Exp Bot 65, 2769–2784.

Dossin, F., Pinheiro, I., Zylicz, J.J., Roensch, J., Collombet, S., Le Saux, A., Chelmicki, T., Attia, M., Kapoor, V., Zhan, Y., Dingli, F., Loew, D., Mercher, T., Dekker, J., and Heard, E. (2020). SPEN integrates transcriptional and epigenetic control of X-inactivation. Nature 578, 455–460.

Dumbliauskas, E., Lechner, E., Jaciubek, M., Berr, A., Pazhouhandeh, M., Alioua, M., Cognat, V., Brukhin, V., Koncz, C., Grossniklaus, U., Molinier, J., and Genschik, P. (2011). The Arabidopsis CUL4-DDB1 complex interacts with MSI1 and is required to maintain MEDEA parental imprinting. EMBO J 30, 731–743.

Fang, X., Wu, Z., Raitskin, O., Webb, K., Voigt, P., Lu, T., Howard, M., and Dean, C. (2020). The 3’ processing of antisense RNAs physically links to chromatin-based transcriptional control. Proc Natl Acad Sci U S A 117, 15316–15321.

Fang, X., Wang, L., Ishikawa, R., Li, Y., Fiedler, M., Liu, F., Calder, G., Rowan, B., Weigel, D., Li, P., and Dean, C. (2019). Arabidopsis FLL2 promotes liquid-liquid phase separation of polyadenylation complexes. Nature 569, 265–269.

Finch-Savage, W.E., and Leubner-Metzger, G. (2006). Seed dormancy and the control of germination. New Phytol 171, 501–523.

Hagai, T., Chen, X., Miragaia, R.J., Rostom, R., Gomes, T., Kunowska, N., Henriksson, J., Park, J.E., Proserpio, V., Donati, G., Bossini-Castillo, L., Vieira Braga, F.A., Naamati, G., Fletcher, J., Stephenson, E., Vegh, P., Trynka, G., Kondova, I., Dennis, M., Haniffa, M., Nourmohammad, A., Lassig, M., and Teichmann, S.A. (2018). Gene expression variability across cells and species shapes innate immunity. Nature 563, 197–202.

Ibarra, S.E., Tognacca, R.S., Dave, A., Graham, I.A., Sanchez, R.A., and Botto, J.F. (2016). Molecular mechanisms underlying the entrance in secondary dormancy of Arabidopsis seeds. Plant Cell Environ 39, 213–221.

Iwafuchi-Doi, M., and Zaret, K.S. (2014). Pioneer transcription factors in cell reprogramming. Genes Dev 28, 2679–2692.

Je, J., Chen, H., Song, C., and Lim, C.O. (2014). Arabidopsis DREB2C modulates ABA biosynthesis during germination. Biochem Biophys Res Commun 452, 91–98.

Kim, D., Paggi, J.M., Park, C., Bennett, C., and Salzberg, S.L. (2019). Graph-based genome alignment and genotyping with HISAT2 and HISAT-genotype. Nat Biotechnol 37, 907–915.

Kushiro, T., Okamoto, M., Nakabayashi, K., Yamagishi, K., Kitamura, S., Asami, T., Hirai, N., Koshiba, T., Kamiya, Y., and Nambara, E. (2004). The Arabidopsis cytochrome P450 CYP707A encodes ABA 8’-hydroxylases: key enzymes in ABA catabolism. EMBO J 23, 1647–1656.

Langmead, B., and Salzberg, S.L. (2012). Fast gapped-read alignment with Bowtie 2. Nat Methods 9, 357–359.

Lee, H.G., Lee, K., and Seo, P.J. (2015). The Arabidopsis MYB96 transcription factor plays a role in seed dormancy. Plant Mol Biol 87, 371–381.

Lee, W.Y., Lee, D., Chung, W.I., and Kwon, C.S. (2009). Arabidopsis ING and Alfin1-like protein families localize to the nucleus and bind to H3K4me3/2 via plant homeodomain fingers. Plant J 58, 511–524.

Lefebvre, V., North, H., Frey, A., Sotta, B., Seo, M., Okamoto, M., Nambara, E., and Marion-Poll, A. (2006). Functional analysis of Arabidopsis NCED6 and NCED9 genes indicates that ABA synthesized in the endosperm is involved in the induction of seed dormancy. Plant J 45, 309–319.

Li, H., Handsaker, B., Wysoker, A., Fennell, T., Ruan, J., Homer, N., Marth, G., Abecasis, G., Durbin, R., and Genome Project Data Processing, S. (2009). The Sequence Alignment/Map format and SAMtools. Bioinformatics 25, 2078–2079.

Liao, Y., Smyth, G.K., and Shi, W. (2014). featureCounts: an efficient general purpose program for assigning sequence reads to genomic features. Bioinformatics 30, 923-

Linkies, A., Graeber, K., Knight, C., and Leubner-Metzger, G. (2010). The evolution of seeds. New Phytol 186, 817–831.

Liu, F., Zhang, H., Ding, L., Soppe, W.J.J., and Xiang, Y. (2020). REVERSAL OF RDO5 1, a Homolog of Rice Seed Dormancy4, Interacts with bHLH57 and Controls ABA Biosynthesis and Seed Dormancy in Arabidopsis. Plant Cell 32, 1933–1948.

Liu, Y., Shi, L., Ye, N., Liu, R., Jia, W., and Zhang, J. (2009). Nitric oxide-induced rapid decrease of abscisic acid concentration is required in breaking seed dormancy in Arabidopsis. New Phytol 183, 1030–1042.

Love, M.I., Huber, W., and Anders, S. (2014). Moderated estimation of fold change and dispersion for RNA-seq data with DESeq2. Genome Biol 15, 550.

Martinez-Andujar, C., Ordiz, M.I., Huang, Z., Nonogaki, M., Beachy, R.N., and Nonogaki, H. (2011). Induction of 9-cis-epoxycarotenoid dioxygenase in Arabidopsis thaliana seeds enhances seed dormancy. Proc Natl Acad Sci U S A 108, 17225–17229.

Molitor, A.M., Bu, Z., Yu, Y., and Shen, W.H. (2014). Arabidopsis AL PHD-PRC1 complexes promote seed germination through H3K4me3-to-H3K27me3 chromatin state switch in repression of seed developmental genes. PLoS Genet 10, e1004091.

Okamoto, M., Kuwahara, A., Seo, M., Kushiro, T., Asami, T., Hirai, N., Kamiya, Y., Koshiba, T., and Nambara, E. (2006). CYP707A1 and CYP707A2, which encode abscisic acid 8’-hydroxylases, are indispensable for proper control of seed dormancy and germination in Arabidopsis. Plant Physiol 141, 97–107.

Pazhouhandeh, M., Molinier, J., Berr, A., and Genschik, P. (2011). MSI4/FVE interacts with CUL4-DDB1 and a PRC2-like complex to control epigenetic regulation of flowering time in Arabidopsis. Proc Natl Acad Sci U S A 108, 3430–3435.

Qi, J., Sun, G., Wang, L., Zhao, C., Hettenhausen, C., Schuman, M.C., Baldwin, I.T., Li, J., Song, J., Liu, Z., Xu, G., Lu, X., and Wu, J. (2016). Oral secretions from Mythimna separata insects specifically induce defence responses in maize as revealed by high-dimensional biological data. Plant Cell Environ 39, 1749–1766.

Questa, J.I., Song, J., Geraldo, N., An, H., and Dean, C. (2016). Arabidopsis transcriptional repressor VAL1 triggers Polycomb silencing at FLC during vernalization. Science 353, 485–488.

Schmitges, F.W., Prusty, A.B., Faty, M., Stutzer, A., Lingaraju, G.M., Aiwazian, J., Sack, R., Hess, D., Li, L., Zhou, S., Bunker, R.D., Wirth, U., Bouwmeester, T., Bauer, A., Ly-Hartig, N., Zhao, K., Chan, H., Gu, J., Gut, H., Fischle, W., Muller, J., and Thoma, N.H. (2011). Histone methylation by PRC2 is inhibited by active chromatin marks. Mol Cell 42, 330–341.

Shevchenko, A., Tomas, H., Havlis, J., Olsen, J.V., and Mann, M. (2006). In-gel digestion for mass spectrometric characterization of proteins and proteomes. Nat Protoc 1, 2856–2860.

Tanaka, M., Kikuchi, A., and Kamada, H. (2008). The Arabidopsis histone deacetylases HDA6 and HDA19 contribute to the repression of embryonic properties after germination. Plant Physiol 146, 149–161.

Vera Alvarez, R., Pongor, L.S., Marino-Ramirez, L., and Landsman, D. (2019). TPMCalculator: one-step software to quantify mRNA abundance of genomic features. Bioinformatics 35, 1960–1962.

Wu, J., Hettenhausen, C., Meldau, S., and Baldwin, I.T. (2007). Herbivory rapidly activates MAPK signaling in attacked and unattacked leaf regions but not between leaves of Nicotiana attenuata. Plant Cell 19, 1096–1122.

Wu, Z., Ietswaart, R., Liu, F., Yang, H., Howard, M., and Dean, C. (2016a). Quantitative regulation of FLC via coordinated transcriptional initiation and elongation. Proc Natl Acad Sci U S A 113, 218–223.

Wu, Z., Zhu, D., Lin, X., Miao, J., Gu, L., Deng, X., Yang, Q., Sun, K., Zhu, D., Cao, X., Tsuge, T., Dean, C., Aoyama, T., Gu, H., and Qu, L.J. (2016b). RNA Binding Proteins RZ-1B and RZ-1C Play Critical Roles in Regulating Pre-mRNA Splicing and Gene Expression during Development in Arabidopsis. Plant Cell 28, 55–73.

Yang, C., Bratzel, F., Hohmann, N., Koch, M., Turck, F., and Calonje, M. (2013). VAL- and AtBMI1-mediated H2Aub initiate the switch from embryonic to postgerminative growth in Arabidopsis. Curr Biol 23, 1324–1329.

Zaret, K.S., and Mango, S.E. (2016). Pioneer transcription factors, chromatin dynamics, and cell fate control. Curr Opin Genet Dev 37, 76–81.

Zhang, X., Clarenz, O., Cokus, S., Bernatavichute, Y.V., Pellegrini, M., Goodrich, J., and Jacobsen, S.E. (2007). Whole-genome analysis of histone H3 lysine 27 trimethylation in Arabidopsis. PLoS Biol 5, e129.

Zhang, Y., Lin, Y.H., Johnson, T.D., Rozek, L.S., and Sartor, M.A. (2014). PePr: a peak-calling prioritization pipeline to identify consistent or differential peaks from replicated ChIP-Seq data. Bioinformatics 30, 2568–2575.

Zhang, Y., Li, Z., Chen, N., Huang, Y., and Huang, S. (2020). Phase separation of Arabidopsis EMB1579 controls transcription, mRNA splicing, and development. PLoS Biol 18, e3000782.

Zhu, D., Mao, F., Tian, Y., Lin, X., Gu, L., Gu, H., Qu, L.J., Wu, Y., and Wu, Z. (2020). The Features and Regulation of Co-transcriptional Splicing in Arabidopsis. Mol Plant 13, 278–294.

Zylicz, J.J., Bousard, A., Zumer, K., Dossin, F., Mohammad, E., da Rocha, S.T., Schwalb, B., Syx, L., Dingli, F., Loew, D., Cramer, P., and Heard, E. (2019). The Implication of Early Chromatin Changes in X Chromosome Inactivation. Cell 176, 182–197 e123.

